# Can category-selective cortex predict categorisation behaviour?

**DOI:** 10.64898/2026.01.23.701404

**Authors:** Timothee Maniquet, Huangxu Fang, N. Apurva Ratan Murty, Hans Op de Beeck

## Abstract

One of the distinctive features of the human visual system is the presence in occipito-temporal cortex (OTC) of regions that show preferential activation to specific categories of visual objects. To understand how this selectivity relates to categorisation behaviour, studies have employed a distance-to-bound approach (DTB), where multivariate brain activity is used to estimate a decision boundary, from which behavioural performance can be predicted. Using this approach, correlations have been found between activity in OTC, and behavioural performance when carrying out certain categorisation tasks. However, it remains unclear what determines where in OTC this correlations can be found, and with which categorisation tasks they can be found. Here, we bridged this gap by relating category-selective regions of OTC, to behavioural performance while participants categorised images as belonging or not to their preferred categories. We adopted a more basic approach and considered simple, univariate activity, rather than relying on decoding to build our DTB. Our results show that activation in regions selective to faces (FFA & OFA), bodies (EBA), and scenes (PPA), is sufficient to predict behavioural performance while categorising images as being faces, bodies, or scenes, respectively. These results are largely consistent across reaction time and motor movements, and generalise to animacy classification. Overall, our data adds to evidence that category-selective regions in OTC can serve to guide categorisation behaviour, and underlines the validity of the DTB approach to address this relationship.

## 1 Introduction

A hallmark of human visual cortex is the presence of regions in occipito-temporal cortex (OTC) that show preferential activation to images of certain categories Bracci and Op de Beeck (2023), for instance faces in the fusiform face area and occipital face area (FFA & OFA Kanwisher, McDermott, and Chun (1997); Puce, Allison, Asgari, Gore, and McCarthy (1996)); bodies in the extrastriate body area (EBA Downing, Jiang, Shuman, and Kanwisher (2001)), and scenes in the parahippocampal place area (PPA Epstein and Kanwisher (1998)). Across these regions, a superordinate selectivity to animate and inanimate objects exists, whereby regions more medial in OTC, including PPA, tend to preferentially respond to images of inanimate objects, whereas regions more lateral, including FFA, tend to preferentially respond to images of animate objects Kiani, Esteky, Mirpour, and Tanaka (2007). This animacy gradient has been described as the progressive preference for living and non-living objects Caramazza and Shelton (1998); Sha et al. (2015); Thorat, Proklova, and Peelen (2019). However, the surface it covers encloses the smaller regions of OTC, and an alternative explanation is that it reflects selectivity to face- and body-like stimuli Leys, Chen, von Leupoldt, Ritchie, and Op de Beeck (2025); Proklova and Goodale (2022); Ritchie et al. (2021).

To understand whether the pattern of response in these regions serves behaviour, research has linked it to categorisation behaviour by estimating a decision criterion directly from brain activity, through a distance-to-bound (DTB) approach Ritchie and Carlson (2016). Making use of this approach, negative correlations were found between the DTB extracted from a classifier trained on animacy-selective regions, and RT in classifying stimuli as animate or inanimate Carlson, Ritchie, Kriegeskorte, Durvasula, and Ma (2014); Grootswagers, Cichy, and Carlson (2018); Grootswagers, Ritchie, Wardle, Heathcote, and Carlson (2017); Ritchie and Op de Beeck (2019); Ritchie, Tovar, and Carlson (2015). Specifically, the animals that are furthest from the neurally derived decision boundary, often mammals, are categorised the fastest in behaviour. Despite the frequent limitation of only finding this correlation in animate stimuli, these studies provide evidence that the organisation of stimulus-evoked activity in visual cortex could be linearly decoded downstream in the brain to guide categorisation behaviour.

These results have been replicated for other categories, including superordinate (e.g. *manmade vs. natural* in Karapetian et al. (2023); Singer, Karapetian, Hebart, and Cichy (2025)) and basic categories (e.g. faces vs. bodies in Grootswagers et al. (2018)). These correlations seem to be found regardless of the task performed during brain recording, even when brain data is recorded while doing a task orthogonal to the categories decoded Carlson et al. (2014); Grootswagers et al. (2018); Karapetian et al. (2023); Ritchie et al. (2015); Singer et al. (2025,?). However, not all categories that can be decoded from brain data show correlations with RT or accuracy. Some classifiers hence perform above chance on brain data, but their distances do not correlate with human categorisation performance (e.g., *human vs. animal* in Grootswagers et al. (2018); living vs. *non-living* in Contini, Goddard, and Wardle (2021)), suggesting that the pattern of stimulus-evoked activity in visual cortex cannot always serve as a direct readout for downstream areas to guide behaviour.

What determines *which categories* can be linked to brain activity, and in turn, *which regions* of OTC can be linked to behavioural performance? Answer this question could hep understand the behavioural relevance of category-selectivity in OTC. A first obvious starting point in this investigation are the categorical distinctions that define the best-known category-selective regions: Faces (vs nonfaces), bodies, and scenes. Strikingly, these distinctions have not been tested yet with the DTB approach.

Here, we asked whether distances to a one-dimensional bound extracted from univariate activity in FFA, OFA, EBA, and PPA can relate to the categorisation behaviour of their preferred categories. While previous fMRI DTB studies looked for correlations DTB extracted from *populations* of voxels, the presence of univariate selectivity provides the opportunity to investigate whether the overall univariate activity would already be sufficient to predict behavioural performance in a categorisation task.

We correlated this univariate DTB with RTs and motor movements in categorising stimuli as belonging to the preferred category of each ROI or not (e.g. “Is this a face? Yes/no” to correlate with FFA). We found that all our ROIs correlated significantly with behavioural performance in categorising images from non-preferred categories. Additionally, FFA and OFA also correlated with performance for images of faces.

Finally, we extended our approach to an animacy classification task, using the same univariate activity which we correlated with the motor movements of participants classifying stimuli as animate or inanimate. Surprisingly, we found that we could largely replicate the patterns of correlations observed with our basic category tasks. This is consistent with an account of animacy as progressive selectivity to human faces and bodies Leys et al. (2025); Proklova and Goodale (2022); Ritchie et al. (2021).

## 2 Methods

### 2.1 Stimulus set

Across all experiments, we used a stimulus set of 185 colour images introduced by Ratan Murty et al (2021 Ratan Murty, Bashivan, Abate, DiCarlo, and Kanwisher (2021). These images include exemplars from one of 4 categories: faces (25 images), bodies (50 images), objects (60 images), and scenes (50 images, see Fig. 1A). Each exemplar was left on its original colour background, and most of the pictures were taken from the THINGS dataset (see Ratan Murty et al. (2021) for more information).

**Figure 1:**
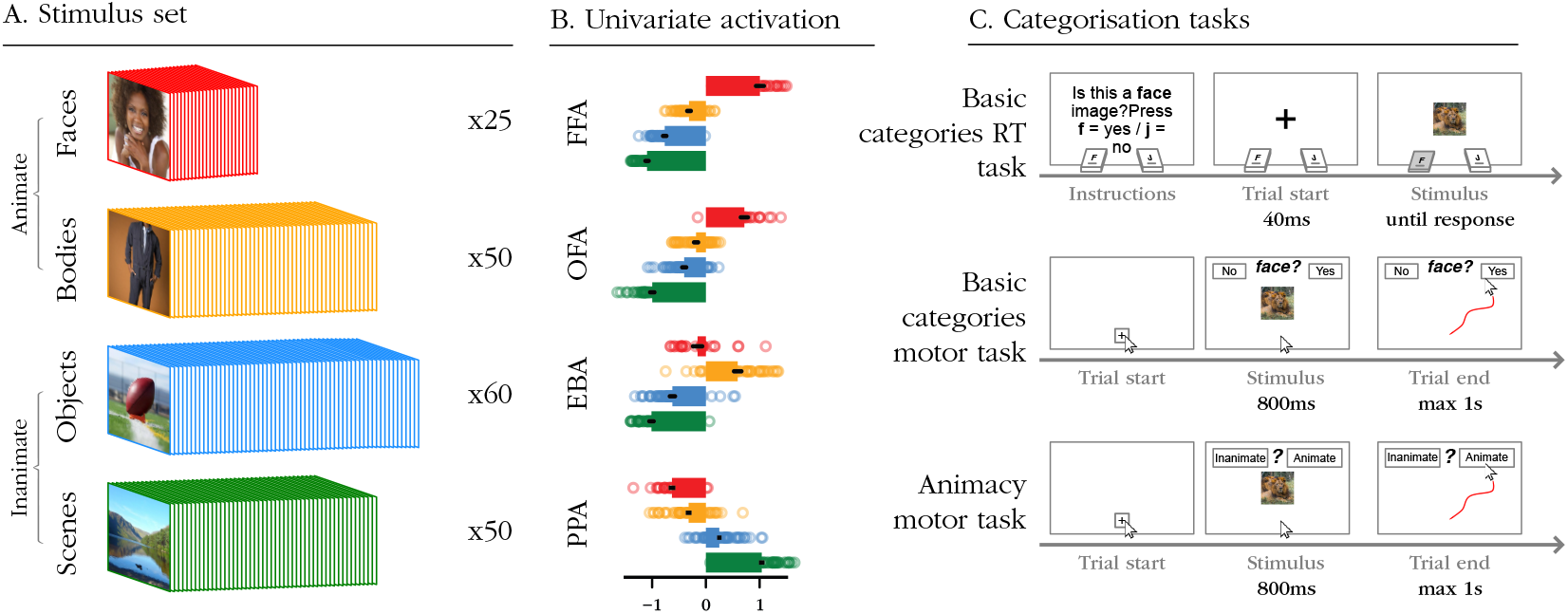
Stimulus set, fMRI data and behavioural tasks. (A) Stimulus set included in the fMRI and all 3 behavioural experiments. 185 images in total, including faces (25 images), bodies (50 images), objects (60 images) and scenes (50 image). Animate images were those of faces and bodies (75 images in total), while inanimate images were those of objects and scenes (110 images in total). (B) Standardised univariate activity per ROI, across categories. Bars indicate average activity for each category of images, error bars indicate SEM. Dots show average activity for individual images. For details on both images and brain data, see Ratan Murty et al. (2021). (C) Categorisation tasks used in this experiment. In both basic categories tasks (RT and motor tasks), participants had to classify images as belonging or not to *faces, bodies, or scenes*. In the animacy task, participants had to indicate whether images were of animate objects or not.

For the three behavioural experiments, we additionally created a smaller, 22-images stimulus set for training. All were freely acessible on the Kaggle dataset website (kaggle.com). The following images were included: 3 images of faces, 6 images of bodies, 6 images of scenes, and 7 images of objects.

### 2.2 fMRI data

All fMRI data used in this project comes from the work of Ratan-Murty and colleagues (see Ratan Murty et al. (2021) for full details). In short, data was collected from 4 participants (2 females) using a 3T MRI scanner, over the scope of 5 scanning sessions during which a localiser was performed, as well as a passive fixation task.

FFA, OFA, EBA and PPA were localised during the first session using a dynamic localiser, consisting of short videos containing five types of stimuli (faces, bodies, scenes, objects, and scrambled objects).

A passive fixation task was repeated over the next four sessions, where each stimulus was repeated at least (20 times across participants. Participants were only instructed to maintain fixation on a fixation cross at the center of each image. Trials consisted in a 300ms presentation of the stimulus, followed by a variable ISI of 3700 to 11700ms. Images were displayed to span 8 dva in size. Univariate activity was then extracted using beta maps from a GLM applied on the preprocessed data (see Fig. 1B for a bar plot visualisation of the univariate activity per ROI).

### 2.3 Basic category RT experiment

#### 2.3.1 Task

For the basic category reaction time experiment, we collected data from 78 participants (69 females, mean age = 19.05 ± 2.25 SD, range 17-31, all of them right-handed). All subjects provided informed consent before participation. The experiment was done onsite at the Faculty of Psychology of KU Leuven, and was approved by the ethics committee of KU Leuven.

Participants performed a keyboard-based categorisation task, where they were asked to categorise as quickly and accurately as possible whether a presented image belonged to a certain category or not (see Fig. 1C). After a short training, they were presented with 18 blocks of the task. During each block, one categorisation task was performed, from one of the three following categories: faces, bodies, and scenes (*is this a face? yes/no, is this a body? yes/no, etc.*). To remove potential motor activity confounds, the keys associated with the yes and no answers (keys F and J) were alternated across blocks. This combination of 3 tasks and 2 key mappings yielded the 6 blocks. All 185 images of the stimulus set were presented in each of the blocks, in random order. The code to run this experiment is publicly available on [GitHub](https://github.com/TimManiquet/RT_exp).

#### 2.3.2 Preprocessing

To ensure good data quality, we first excluded participants whose average accuracy was below 0.7 (5 participants excluded). We then ensured all trials had reasonable RT values and were in the range of the 0.5th to the 99.5th percentiles (from 0.17 to 1.93 seconds, 2376 trials excluded). Finally, given the long nature of the task, we calculated average accuracies per participant and per block, and removed blocks with average accuracies below 0.6 (30 blocks removed). After preprocessing, 73 participants were left, for a total of 235164 trials.

### 2.4 Basic category motor movement experiment

#### 2.4.1 Task

After collecting keyboard-based RT data on the basic categories task, we replicated the paradigm with mouse tracking to collect time-resolved motor data (see Fig. 1C). Instead of responding to images by pressing a key on a keyboard, participants here responded by moving their mouse cursor towards a *yes* or a *no*button, positioned at the top-left and top-right of the display.

We designed our mouse tracking experiment broadly following the procedure described by Koenig-Roberts and colleagues Koenig-Robert, Quek, Grootswagers, and Varlet (2023). To ensure a comparable movement trajectory, each trial started when participants pressed a small *start* button at the bottom center of the display. At that point, images appeared on screen for 800 ms. Instructions required to move as quickly as possible towards the correct answer. To ensure fast movement onsets, trial durations were capped to one second, at which point the task would move on to the starting box of the next trial.

The full experiment consisted in 4 blocks, one for each of the 4 categories in the stimulus set: faces, bodies, objects and scenes (*is this a face? yes/no, is this a body?* yes/no, etc.). The order of the blocks was randomised. Within each block, all 185 images were presented once, in random order. Button positions were switched halfway through the block after a short break to avoid potential motor activity confounds.

Data was collected from a total of 81 participants (59 females, mean age = 19.78 ± 4.82 SD, range 17-56, 76 right-handed). The experiment was conducted online and was approved by the ethics committee of KU Leuven. Participants signed a digital consent form before starting the task. Code for this experiment is available on [GitLab](https://gitlab.pavlovia.org/hoplab/mouse_tracker_category).

#### 2.4.2 Preprocessing

We took the following steps to preprocess the mouse trajectory data: trials were aligned by flipping the x coordinates when necessary, so that all trials could be analysed assuming the No button was on the left and the Yes button on the right of the display. All x and y data points were scaled and standardised according to individual screen sizes to bring all trials into a common space, spanning from −1 to 1 for the x axis (with the starting box roughly at 0, the No box roughly at −1, and the Yes box roughly at 1), and from 0 to 1 for the y axis.

Given the challenging nature of the task (in particular due to the capping of trial duration to 1s), and to try and capture with maximum sensitivity the motor movement dynamics associated with each image, we adopted exclusion criteria as light as possible. First, we excluded participants who had completed (i.e. managed to click on a response button) less than (200 out of the possible 555 total trials of the task (4 participants excluded).

We then excluded outlier trials where the trajectory significantly differed from normal. For that we looked at trajectories that did not move far enough vertically (end point less than 0.05 away from starting box); trajectories too short (that generated less than 10 coordinate points, compared to an average of 33); and trajectories that ended below the starting box (18275 trials excluded). After preprocessing, 77 participants were left, for a total of 41665 trials.

### 2.5 Animacy motor movement experiment

#### 2.5.1 Task

We adapted the motor movement paradigm we created for basic categories to turn it into an animacy classification task (see Fig. 1C). Methodological details for this task were largely in line with those of the basic category motor movement experiment, with the exception that the animacy task was this time repeated 3 times across blocks (the same division was adopted within blocks to switch response buttons). Instructions required participants to classify images as *animate* if they portrayed living things, and as *inanimate* if they portrayed non-living things.

Data was collected from a total of 84 participants (82 females, mean age = 18.26 ± 1.04 SD, range 17-24, 77 right-handed). The experiment was conducted online and was approved by the ethics committee of KU Leuven. Participants signed a digital consent form before starting the task. Code for this experiment is available on [GitLab](https://gitlab.pavlovia.org/hoplab/tim_mouse_tracker_test).

#### 2.5.2 Preprocessing

Preprocessing for the animacy motor movement data followed the same steps exposed in 2.4.2. After preprocessing, 75 participants remained, 16546 trials were excluded for a total of 30074 trials left.

## 3 Results

### 3.1 Univariate activity from category-selective cortex correlates with RT in categorising basic categories

In our RT behavioural experiment, participants were instructed to categorise images as belonging, or not, to one of three basic categories: faces, bodies, and scenes. Here, we tried to understand whether activation in patches of cortex selective to these categories could guide categorisation behaviour. To this end, we correlated univariate brain activity and behavioural RTs, and compared the correlations to predictions from a DTB approach.

We averaged univariate activity from FFA, OFA, EBA, and PPA, per image, across participants and hemispheres. We averaged RTs from accurate trials only from participants in the basic category RT experiment (n=73, see 2.3), per image and per task (one task per target category: faces, bodies, and scenes, see Fig. 2A). We then correlated average brain activity to average RT, for each ROI with the RTs to the categorisation task of its preferred category (FFA & OFA: face-, EBA: body-, PPA: scene categorisation task, see Fig. 2B).

**Figure 2:**
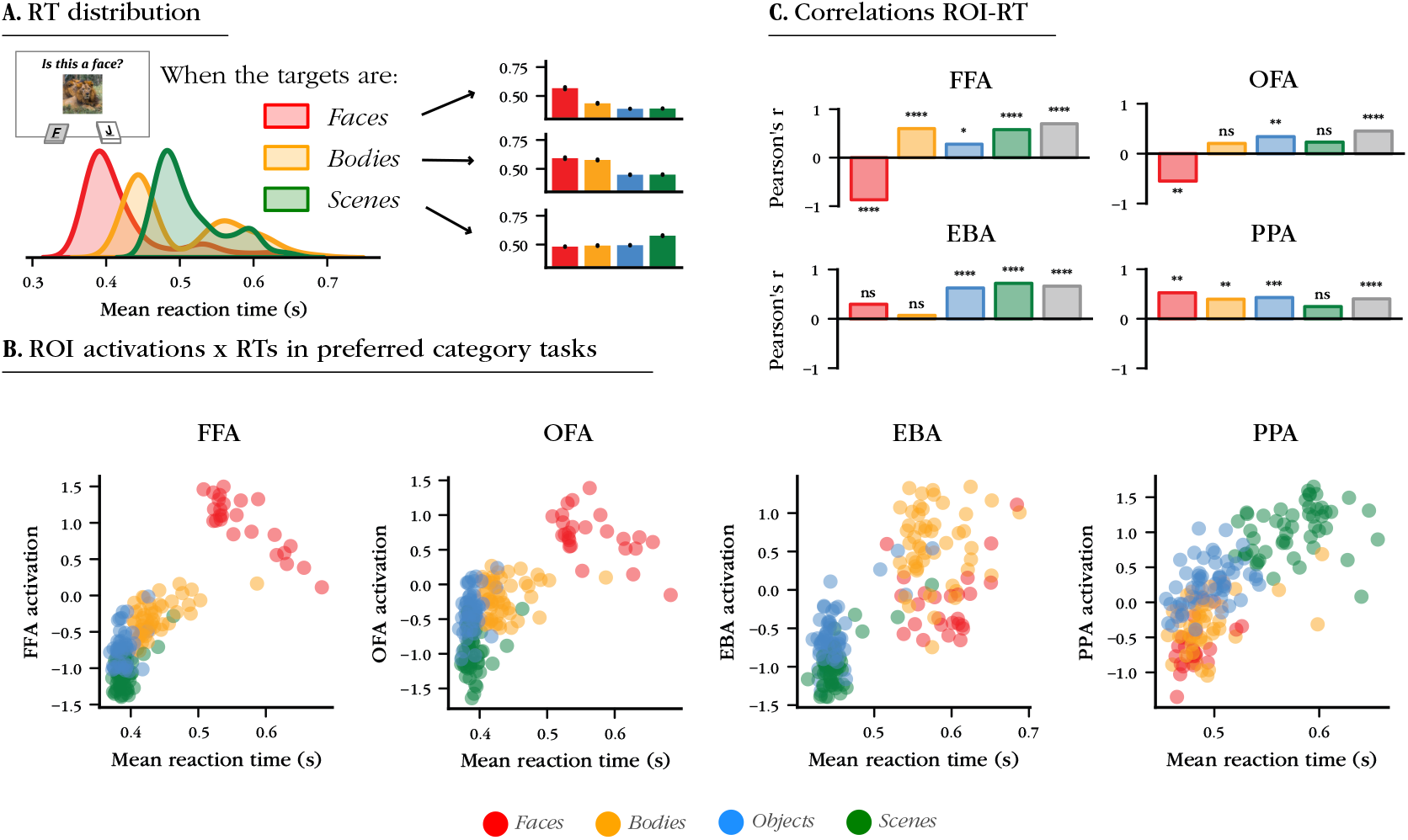
Results from the basic category RT experiment correlate with ROI univariate activity. (A) Distribution of RTs in each of the three tasks, using only correct trials. For each task, participant had to indicate whether the image belonged to the target category or not, using keys on a keyboard. (B) Correlations between average RTs per image and average univariate activity per image. Each plot shows data from the task where the preferred category of the ROI was the target (FFA activation is shown against RTs in categorising faces, EBA activation against RTs in categorising bodies, etc.). (C) Correlation scores from the data in (B). For each ROI, we correlated the RTs and univariate activity per image. Results are split between categories. Grey bars indicate the correlation between univariate activity and RTs for stimuli from all non-preferred categories pooled together (e.g. all images of bodies, objects, and scenes for FFA and OFA).

We expected activations from each ROI to be negatively correlated with RTs to stimuli from its preferred category, as would be predicted from a DTB approach. We found such a negative correlation for both of our face-selective regions (FFA: r=-0.87, OFA: r-0.55; both p<0.0001 - all correlations tested with two-sided t-tests), but found a null correlation for EBA (r=0.07, p=0.63), and a non-significant *positive* correlation for PPA (r=0.24, p=0.09; see Fig. 2C).

Then, we expected activations to positively correlate with RTs to stimuli from non-preferred categories, as indicative of longer RTs to images closer to the boundary defined by univariate brain activity. We found this significant positive correlation for all our ROIs (FFA: r=0.7; OFA: r=0.45; EBA: r=0.67; PPA: r=0.4; all p<0.0001). Overall, these results indicate that univariate activity from category-selective patches of cortex can be linked to RTs from categorisation tasks.

### 3.2 RT correlations with univariate activity from category-selective regions replicate in motor movements

We next replicated our analyses, using time-resolved behavioural data in place of RTs, considering RTs alone do not capture the entire decision process behind categorisation. We adopted a more sophisticated approach to ensure the pattern of results would not differ significantly if we were able to look at image decisions before decisions were actually made.

To this end, we designed a mouse-tracking experiment, where participants categorised images as belonging, or not, to the same basic categories as before (faces, bodies, scenes), but this time responded by clicking a button on the screen rather than a key on their keyboard. In doing so, we were able to analyse the trajectory of their mouse cursor, and take that motor movement as a proxy for the different steps taken in making a categorisation decision.

We focused on the horizontal (x) position of the mouse cursor across time, as it reflects the proximity of participants to making either a *yes* or *no* decision at every time point during a trial. We extracted horizontal position at each time point, separately for each category task (see Fig. 3A), and averaged it over participants to obtain one time-resolved average horizontal trajectory per image (see Fig. 3B).

**Figure 3:**
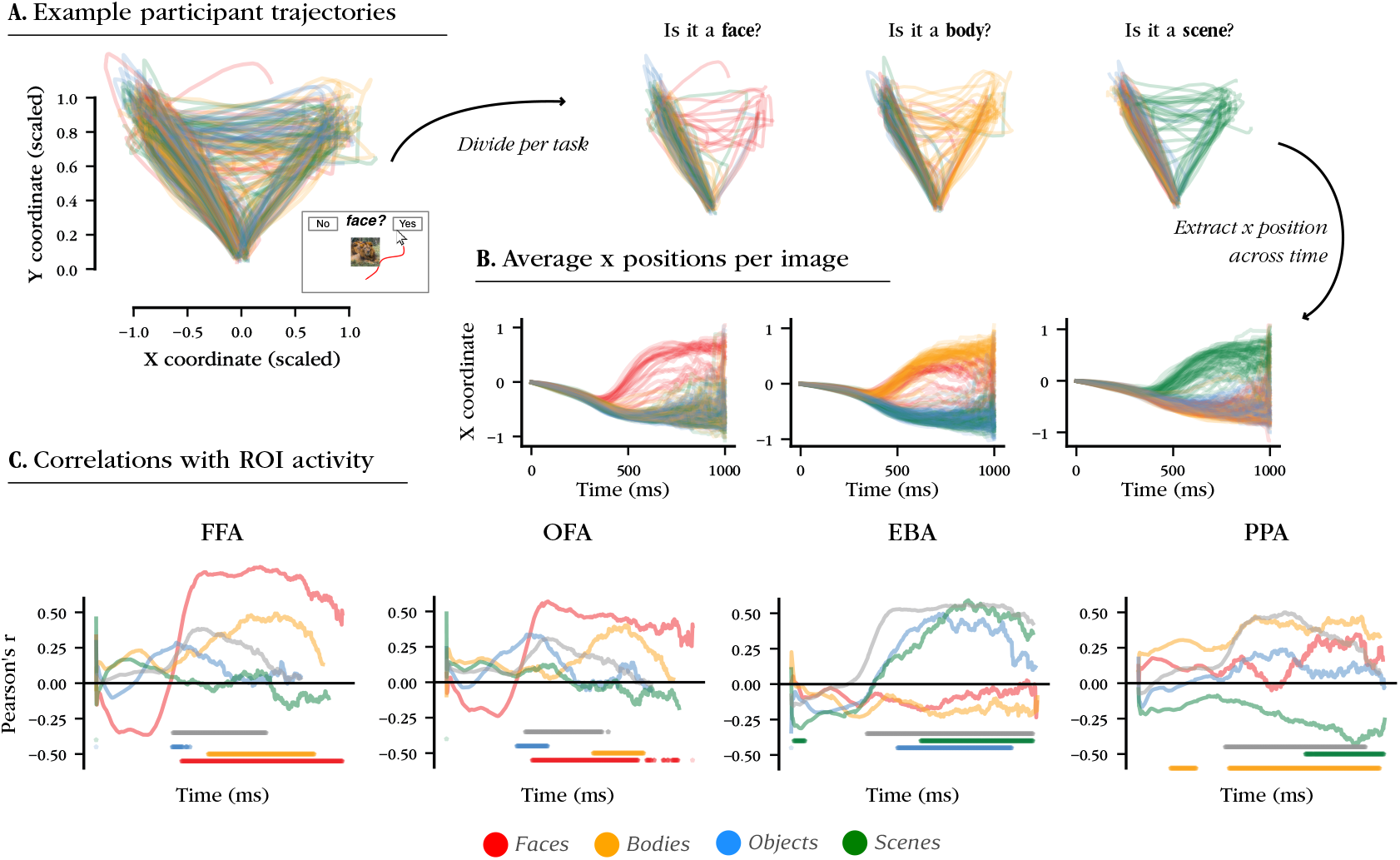
Results from the basic category motor task largely replicate those from the RT task. (A) All trajectories (one per trial) from one example participant. Trajectories are flipped when necessary so that the yes button is always on the right (close to x=1.0). The left side plot shows all trajectories from one experimental session, i.e. pooled across tasks. Dividing trials based on the target category, we get the plots on the right side (e.g. all trials from that example participant for when the target was a face under the *Is this a face*? title). (B) Average horizontal (x) position per image and per task, across time. Average data from all participants is shown here. The y axis represents horizontal position, with more positive values closer to the *yes* button. The x axis represents time (with trials capped at 1 second). (C) Correlations between average x position per image and average ROI activation per image, across time. As in Fig. 2B, each ROI is shown against data from trials where its preferred category was the target. Dots below the graphs indicate time points where the correlation is significantly different from zero. Correlations are divided across images from each category (one line per category). The grey line shows correlations for all stimuli from non-preferred categories pooled together.

From the horizontal position of each image across time, we replicated the correlation analysis shown in Fig. 2B-C, this time calculating one correlation value per time point (1000 values in total, for trials of 1 second). Just as in 3.1, we correlated activations from each ROI to horizontal positions obtained from its preferred category task.

We made similar predictions as per a DTB approach, with the one difference that for preferred images, positive correlations were expected: what were previously short RTs would now be larger positive horizontal values (closer to 1.0, i.e. the *yes* button), hence a positive correlation is to be expected between their horizontal position and ROI activity. We still predicted a positive correlation for non-preferred images, as images yielding lower activity in a given ROI should also be lower on the horizontal position (closer to −1.0, i.e. the no button).

We first looked for positive correlations between ROI activations and horizontal position for preferred category images (see Fig. 3C). Firstly, both our face-selective ROIs showed strong correlations with face images. FFA had its peak correlation (r=0.82) at 659ms, with values significantly larger than 0 (p<0.05) from 349ms onwards. OFA had its peak correlation (r=0.61) at 383ms, with values significantly larger than 0 (p<0.05) from 339ms onwards. Secondly, we did not find a significant correlation between EBA activation and the horizontal position of body images. Thirdly, and contrary to predictions, we found *negative* correlations between PPA activation and horizontal positions of scene images, with peak value (−0.44) at 870ms (significantly below 0 from 667ms onwards).

We then looked for positive correlations between ROI activations and horizontal position for non-preferred category images, with the same assumption that higher activation would associate to images closer to the boundary, and hence to images further away from the *no* response button. We were able to find such a correlation for all our ROIs (peak values: 0.38 for FFA, 0.3 for OFA, 0.57 for EBA, and 0.5 for PPA).

Strikingly, when looking at results from the basic category motor task, we found that they largely replicate results from the basic category RT task. In place of finding temporal dissociations in the time-resolved signal which would bring nuance to the previous section, we found the almost exact same patterns of correlation with our four ROIs. (1) First, where we found a negative correlation between univariate activity in FFA and OFA and RT to face images, we found a corresponding positive correlation with horizontal positions for face images (Fig. 3C FFA and OFA, red lines). (2) Second, where we found a positive correlation between univariate activity for all our ROIs and RT to non-preferred images, we found a corresponding positive correlation with horizontal positions for non-preferred images (Fig. 3C all four plots, grey lines). And (3) third, where we did not find the expected negative correlations between univariate activity in EBA and RT to body images, we also did not find the corresponding positive correlation with horizontal position for body images (Fig. 3C EBA, yellow line). The only exception to this pattern of replication is in the unexpected negative correlation that was found late on between univariate activity in PPA and horizontal position for scene images (we come back to this in section 3.4).

This conjunction across our ROIs of the correlations found with simple RTs and with motor movements consolidates our results, and suggest that motor movements can reliably capture the relationship between categorisation performance and brain activation.

### 3.3 Univariate activity in category-selective regions correlates with animacy categorisation behaviour

We finally extended our approach to the categorisation of animacy. Several previous publications have already found correlations between RTs in categorising objects as animate or inanimate, and multivariate activity in OTC (Carlson et al. (2014); Ritchie and Op de Beeck (2019); Ritchie et al. (2015)). On the other hand, it has been suggested that the animacy continuum observed in OTC is driven by the progressive selectivity to face-like and body-like features (Leys et al. (2025); Proklova and Goodale (2022); Ritchie et al. (2021)). We hence wondered if we could extend our correlations to an animacy categorisation task. Therefore, we asked whether univariate activity in patches of cortex with selectivity for basic categories (e.g. faces and bodies) could also be linearly readout by the brain to guide behaviour in categorising the superordinate categories of animate and inanimate.

Considering the results presented above (see 3.2), we chose to use a motor movement task to investigate animacy, as it is able to reproduce results found with simple RTs and provides an extra time-resolved sensitivity. We therefore designed a task where participants classified images as animate or inanimate, and used the mouse cursor movements to create average x positions across time per image, which we then correlated to brain activations (see Fig. 4A-B).

**Figure 4:**
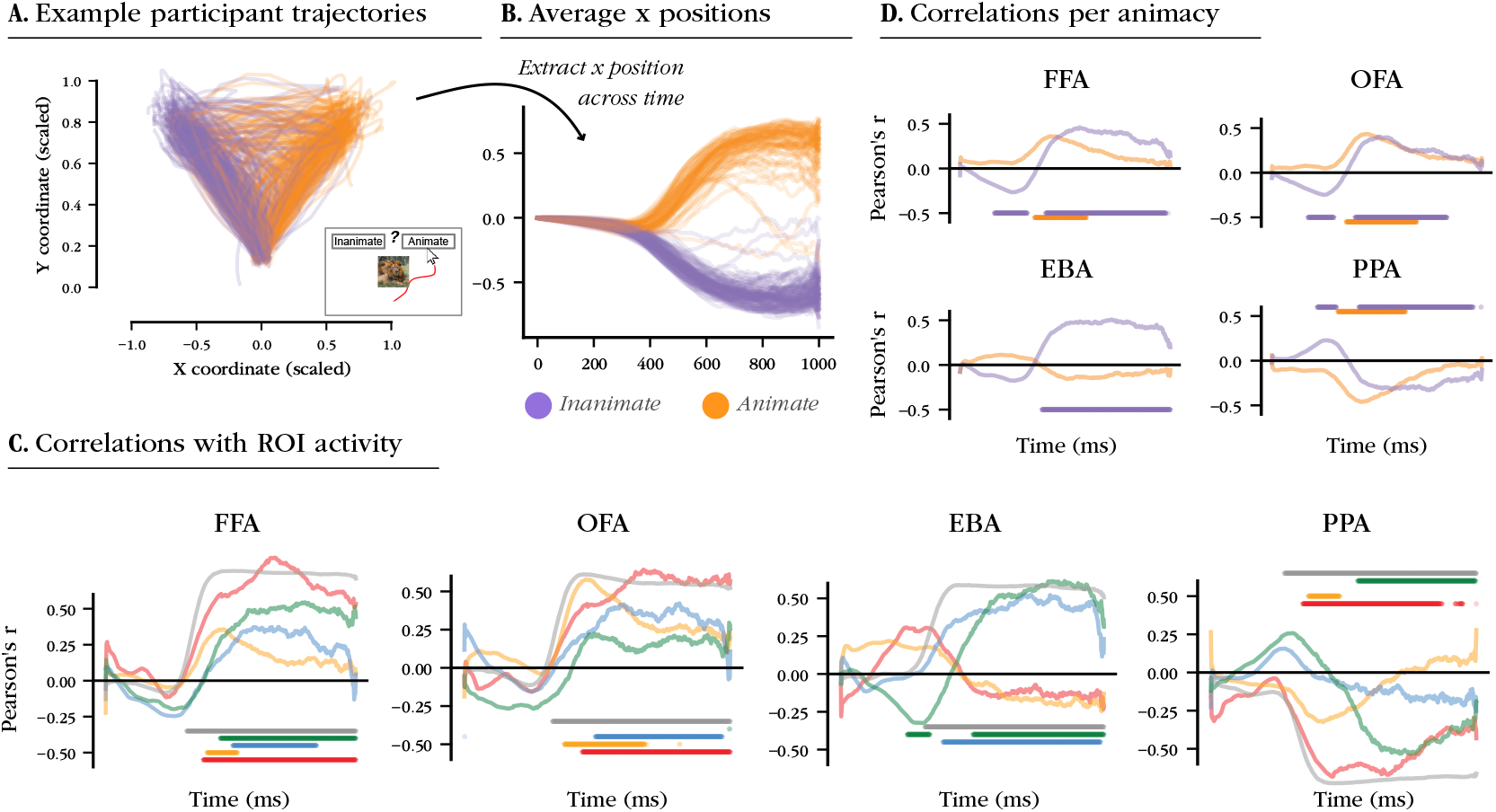
Conjunction of animacy motor task and basic category motor task results. (A) All trajectories (one trajectory per trial) for one example participant. Trajectories were flipped when needed to align the *animate* response button to the right side of the screen, close to x=1. (B) Average horizontal position per image, for all participants. Positions more positive are closer to the *animate* button. (C) Correlations between ROI univariate activity and average horizontal position per image across time. As in Fig. 3B, correlations are divided per categories, with the grey line representing images from all the non-preferred categories. Dots below or above the plots indicate time points where correlations are significantly different from zero. (D) Correlations between ROI univariate activity and image horizontal position across time, pooled between images of animate objects (faces and bodies) and inanimate objects (objects and scenes).

Just like previously, we expected to find correlations in line with a DTB interpretation. Here we have to consider what we observed in the basic category task (taking the motor movement version as the comparison). In that task, we found all expected correlations in FFA and OFA, so for these ROIs the predictions in the animacy categorisation are straightforward. Instead, for EBA and PPA, we only found the predicted (positive) correlation for non-preferred categories, again resulting in straightforward predictions here for those images. However, EBA showed no correlation and PPA an opposite correlation for the preferred category, which should result in similarly non-straightforward predictions in animacy categorisation – at least if our assumption is correct that category-selective ROIs can predict animacy categorisation RTs the same way they can predict face, body, and scene categorisation RTs.

The straightforward predictions are that activations from regions selective to animate categories (faces, bodies, i.e. FFA, OFA, and EBA) should positively correlate with horizontal positions (stimuli eliciting more activity in those regions closer to the *animate* button). Conversely, we expected activations regions selective to inanimate categories (scenes, i.e. PPA) to correlate *negatively* with horizontal positions. This is because trajectories from the animacy categorisation task were aligned so that the *inanimate* button would be on the left, as a result of which more inanimate images are expected to fall closer to −1, and animate images closer to 1.

Results matched these straightforward predictions for the face-selective regions. FFA and OFA showed strong, positive correlations with face images (FFA: peak correlation r=0.85 at 674ms, significantly above 0 from 389ms onwards; OFA: peak correlation r=0.64 at 673ms, significantly above 0 from 443ms onwards), as well as with all non-preferred images (FFA: peak correlation r=0.76 at 504ms, significantly above 0 from 323ms onwards; OFA: peak correlation r=0.61 at 457ms, significantly above zero from 331ms onwards, see Fig. 4B). These basic category results are reflected at the superordinate level: when aggregating bodies and faces together as animate categories, we found significant positive correlations of FFA and OFA activations with both animate and inanimate images (see Fig. 4C)

We did not find a positive correlation between EBA activity and body images in the animacy categorisation task, in line with the absence of a correlation for preferred images in the basic category task. Conversely, we found a positive correlation between EBA activity and non-preferred images (peak correlation r=0.59 at 466ms, correlation significantly above 0 from 319ms onwards). This also translated into no correlation with animate images, and a significantly positive correlation with inanimate images.

Lastly, we found a significant, negative correlation between PPA activation and horizontal position for scene images in the animacy categorisation task (peak correlation: r= −0.54 at 750ms, significantly below 0 from 552ms onwards). This correlation, in line with our expectations, contradicts what was found in the basic category RT task and the basic category motor task. We also found a significant negative correlation between PPA activation and horizontal position for non-preferred images (peak correlation r=-0.73 at 461ms, significantly below 0 from 277 ms onwards). This in turn translated into a significant negative correlation with both animate and inanimate images when pooled at the superordinate level.

To sum up, when looking at results from the animacy motor task, we again found correlations with univariate ROI activity that replicate those from the basic category motor task. Namely, we found that (1) the straightforward prediction is true for OFA and FFA for face images; (2) the straightforward prediction is true for all ROIs for non-preferred images, and (3) the straightforward prediction is not verified for EBA univariate activity and horizontal position for body images. The only major divergence from patterns of results in 3.2 is that of PPA, for which the straightforward prediction was found to be true for scene images.

In sum, overall, the brain-behaviour correlations in the animacy motor task show a close correspondence to brain-behaviour correspondences in the basic category motor tasks.

### 3.4 Category membership ratings correlate with behaviour more than univariate activity

To try and explain what is driving these results we analysed category membership ratings collected by Ratan-Murty and colleagues Ratan Murty et al. (2021). Those ratings were collected by asking participants to align images on the screen based on how face-, body-, or scene-like they were. These category likeness ratings were collected online on 115 participants, with a procedure that allowed for a measure of reliability, so that only the more reliable participants were kept in the final analyses (n=106 for faces, n=115 for bodies, n=60 for scenes).

We first asked whether the rated category membership of an image is predictive of categorisation performance for that image. We used RTs from the basic category keyboard-based experiment (see 2.3) as a benchmark of categorisation performance for our images, and correlated it with category membership ratings.

In line with the DTB approach used before, we expected that higher ratings on a target category would negatively correlate with RTs in categorising it. For instance, if a face was rated as highly face-like, then we predicted that it would be associated with a *faster* RT. In that same example, conversely, if a non-face stimulus (e.g. a scene image) was rated as highly face-like, then we predicted that it would be associated with a *slower*RT.

We found this prediction to be true of all categorisation tasks for ratings of non-target images (see Fig. 5A). This means that for all tasks, the ratings of non-target images on their membership to the target category is correlated with RT in categorising them as non-targets. For target images, only ratings of face-likeness and scene-likeness correlated with RTs in categorising faces and scenes, respectively, while no correlation was found for bodies.

**Figure 5:**
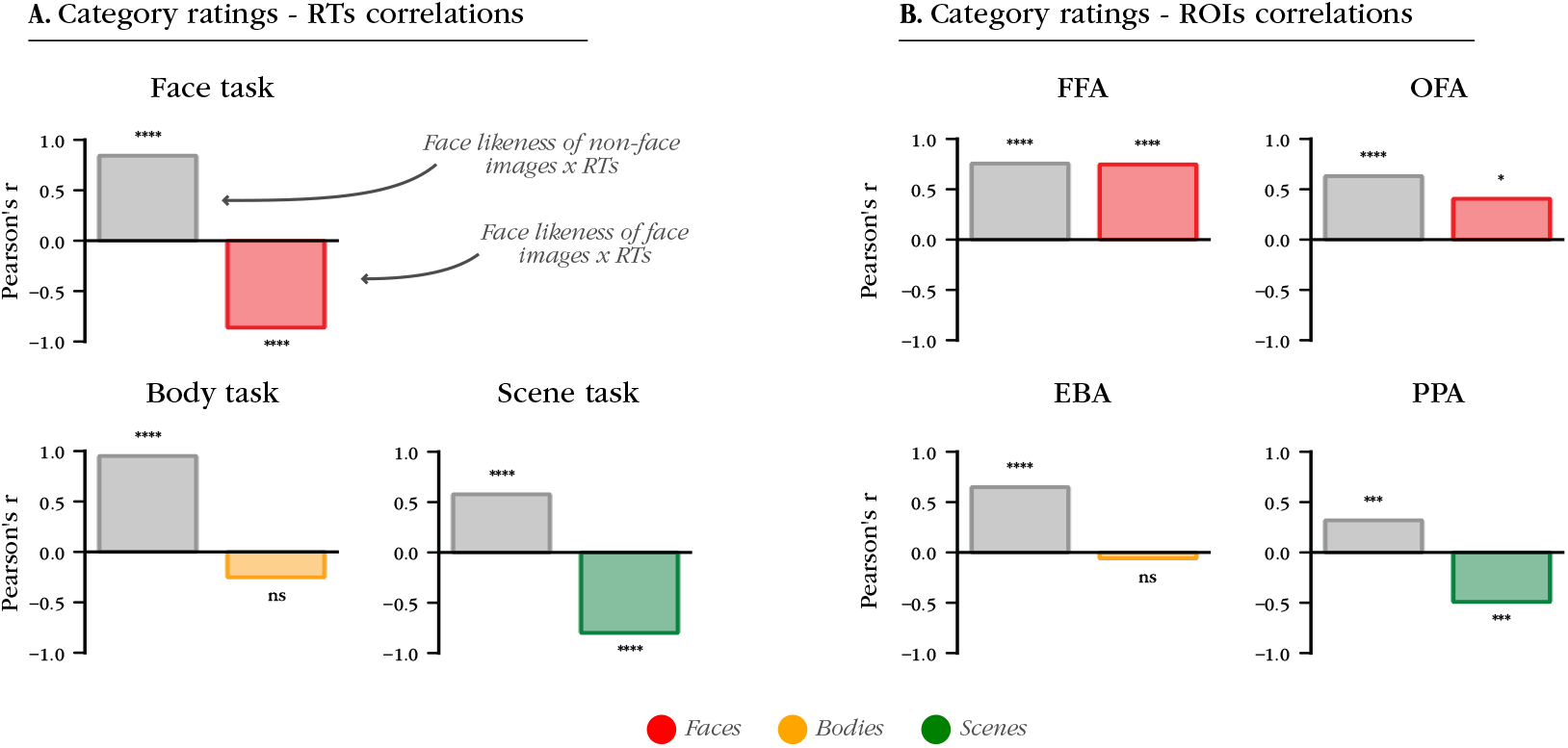
Category membership ratings explain most correlations. (A) Correlations between category membership ratings and average RTs in the basic category RT task. For each task, RTs are correlated with ratings from the target category during that task (e.g. RTs from the task *Is this a face? yes/no* are correlated with ratings on face likeness). Colour bars indicate correlations between ratings for the target categories and RTs to images from the target categories. Grey bars indicate correlations between ratings for the target categories and RTs to images from all nontarget images. (B) Correlations between ROI univariate activity and category membership ratings. Just like in (A), ratings for the likeness to the preferred category of each ROI is correlated with activations from that ROI (e.g. ratings of face likeness are correlated with activations from FFA). Grey bars indicate correlations with the category likeness of stimuli from non-preferred categories.

We then applied this approach to ROIs, and correlated brain activations with category membership ratings. We expected positive correlations: if brain and behaviour aligned, then face likeness, for instance, should positively correlate with activations in FFA. We looked for these positive correlations in preferred categories (faces for OFA and FFA, bodies for EBA, and scenes for PPA), and in non-preferred categories pooled together. Among preferred categories, only face-selective regions were found to correlate positively with face likeness ratings for faces, with EBA not correlating significantly with body likeness ratings for bodies, and PPA correlating significantly negatively with scene likeness for scenes. Note that these unexpected correlations of EBA and PPA are in line with results from previous section (see 3.1, 3.2, 3.3).

Overall, these results show that category membership ratings relate differently to behaviour and ROI activation. On the one hand, categorisation RTs for faces and scenes seem to be linearly correlated with face likeness and scene likeness ratings, respectively. This indicates that, for these two categories, how much participants judge images as target-like directly maps onto the speed of their perceptual decision. This does not apply for bodies, for which body likeness judgements seem orthogonal to perceptual decision speed.

On the other hand, face-selective regions show a similar pattern to face-selective categorisation tasks, whereby face likeness ratings directly map onto ROI activations for FFA and OFA. This is not true of EBA, which shows a non-significant correlation to body-likeness for body stimuli, and of PPA, which shows a negative correlation to scene-likeness for scene stimuli.

Taken together, results on preferred category images paint a different picture across ROIs. First, they point to FFA and OFA as particularly suitable for a direct readout by other parts of the brain for guiding categorisation behaviour, which is probably mediated by a pattern of univariate activations very close to the pattern of face likeness ratings of images. EBA, on the other hand, appears to respond to body images in a pattern orthogonal to body likeness ratings, and therefore, orthogonal to body categorisation behaviour. Finally, PPA appears to respond to scene images in a manner opposite to that of behaviour, with weaker responses to stimuli that would be rated high on scene likeness, and therefore, would be faster to classify as scenes.

## Discussion

In this work, we asked whether univariate activity in FFA, OFA, EBA, and PPA, can drive categorisation behaviour. We build on previous DTB studies with four main innovations:

- We used and compared category-selective ROIs, (instead of OTC and/or early visual cortex more broadly Carlson et al. (2014); Grootswagers et al. (2018); Ritchie and Op de Beeck (2019), or a single of those regions Singer et al. (2025));
- Our stimulus set was directly designed to elicit activations in these regions Ratan Murty et al. (2021), and was used in both a basic and a superordinate categorisation task;
- We calculated a one-dimensional DTB through univariate activity directly;
- We combine two behavioural metrics: RTs and time-resolved mouse tracking-based motor movements, allowing us to measure categorisation behaviour over time.

### 4.1 DTB predictions are partially but consistently found across experiments for category-selective ROIs

Across our three behavioural experiments, we made two types of hypotheses. Type *a*: we expected activation amplitudes to predict easier categorisation of images from preferred categories. For instance, if FFA had a high response amplitude to a certain image of a face, then we expected this image to be easily categorised as a face. Type *b*: we expected activation amplitudes to predict harder categorisation of images from non-preferred categories. In the same example, an image of a scene that elicited a high FFA activation was expected to be harder to categorise as a non-face. In the first experiment, regarding hypothesis *a*, we found a negative correlation between RTs and univariate activation only for FFA and OFA, with EBA and PPA not correlating with body and scene stimuli, respectively. Conversely, regarding hypothesis *b*, we found significant, positive correlations between all ROI activations and RTs in categorising stimuli from non-preferred categories.

The second experiment replicated results from the first experiment, using horizontal position from a motor movement task instead of RTs. The only exception was PPA, with a late, significant negative correlation with performance (when a positive one would have been expected, had hypothesis *a* been true).

The realisation of hypothesis b across these two experiments and all ROIs shows that univariate activity *can* be *sufficient* to correlate with categorisation performance. It also corroborates the long-standing observation that patches of category-selective cortex, despite showing a preference for a given category, also respond in a meaningful way to stimuli from other categories Haxby et al. (2001).

The realisation of hypothesis *a* in FFA and OFA is largely in line with literature connecting the face network to face-recognition behaviour (see e.g. Pitcher, Walsh, Yovel, and Duchaine (2007)). Additionally, results from category membership ratings show that FFA especially, but also OFA, largely agree with human participants in judging which images are more or less face-like (see Fig. 5B).

The non-realisation of hypothesis a in PPA aligns with *a* recent report showing no correlation between PPA and RT in categorising scene images Singer et al. (2025). Additionally, category membership ratings (see Fig. 5B) seem to address the unexpected negative correlation of PPA in our second experiment (see Fig. 3C): while in line with participant judgement, they are actually *inverse* to PPA response (see Fig. 5). This could be explained by the rectilinearity of images: while PPA tunes more strongly to rectilinear image contents (manmade and interior scenes, see

Fig. S8)Kravitz, Peng, and Baker (2011); Nasr, Echavarria, and Tootell (2014), participants judged natural scenes more scene-like, and categorised them faster than manmade scenes (see RTs for scene images in RT scene categorisation task in Fig. S11, as well as larger area under the curve for scene images in motor scene categorisation task in Fig. S14).

The non-realisation of hypothesis *a* in EBA, lastly, seems to indicate that EBA activity did not map directly onto body classification behaviour for body images, despite reports that EBA is linked to body recognition performance Pitcher, Charles, Devlin, Walsh, and Duchaine (2009). This might be because our stimuli lack variability in the body features that EBA is specifically tuned (see *limitations* 4.4). Regardless, EBA correlated with performance for other categories of images, showing that it does tune to image features that are relevant to categorisation behaviour.

### 4.2 Animacy behaviour is recapitulated in basic category behaviour

In our third experiment, we asked whether univariate activity amplitude in our ROIs would also correlate with behaviour in classifying images as animate or inanimate. We were motivated by research suggesting that the animacy continuum in OTC is the result of progressive selectivity to human face- and body-likeness, rather than a taxonomy-like continuum Leys et al. (2025); Proklova and Goodale (2022); Ritchie et al. (2021). If true, this account would predict that animacy categorisation behaviour would be largely similar to face and body categorisation behaviour.

Interestingly, we largely replicated what we found in the basic categorisation task experiments (see Fig. 4) regarding our hypotheses *a* and *b*, with the one exception of PPA, which was found to be in line with hypothesis a), in contradiction with results found in our first two experiments. Overall, our results are in line with accounts of animacy selectivity as the combination of selectivities to various basic categories Leys et al. (2025); Proklova and Goodale (2022); Ritchie et al. (2021). Additionally, this account would predict similar correlation trajectories across basic and animacy categorisations, which is in line with what we found (e.g. first significant correlation with non-preferred stimuli, average across all 4 ROIs: 328ms for basic categorisation tasks, versus 327ms for animacy categorisation task, Figs. 3C vs. 4C).

### 4.3 Interpretation of our results in light of previous DTB studies

Several studies until now have found correlations with behavioural performance only for animate objects. We did not replicate this effect (see 3.3). We found correlations with inanimate stimuli both in the animacy (see Fig. 4D) and basic categorisation tasks (see FFA and OFA with scene and objects in 2, for instance). Overall, the animacy status of our stimuli did not seem to drive whether they would correlate with ROI activation.

This animacy asymmetry has previously been explained by the absence of a real *inanimate* dimension in the brain Carlson et al. (2014), or the use of a strategy of *A* - not A whereby paying attention to features of category A alone is enough Grootswagers et al. (2018). The present correlations for inanimate stimuli during animacy categorisation, combined with the similarity of those correlations in all basic categorisation tasks that explicitly posed a binary choice to participant (e.g. *face - not face*) suggests a different interpretation. The presence or absence of correlation might be more readily explained by stimulus sampling, and by the brain region that stimuli are correlated with.

Task behaviour at the time of brain recording could also influence our results Ritchie, Wardle, Vaziri-Pashkam, Kravitz, and Baker (2025). For instance, EBA could have not correlated with the categorisation of body stimuli because subjects did not need to attend to body-like features while being scanned. If this is true, the fact that some brain-behaviour correlations are found in the absence of any task demand suggests that activation in certain regions of OTC automatically aligns with behaviourally-relevant dimensions of stimuli. FFA provides a good example of such a region, as its activation directly maps onto face categorisation behaviour, by showing sensitivity to the same features that participants judge as face-like (see 3.4).

### 4.4 Limitations of this study

Our study has some limitations. Firstly, we used a limited number of stimuli, potentially not allowing us to sample across a large variety of category features (e.g. the lack of effect with body stimuli could be due to too little variability in body image features).

Secondly, a disadvantage of our DTB approach is that we apply binary category labels onto our stimulus set. While the continuous nature of ROI activation and behavioural metrics (RT and mouse position) allow us to tap into the continuous nature of category selectivity, we measured correlations across images grouped together on the basis of a binary category label. These labels are a useful approximation, as confirmed by the correlations we describe in this project. However, using categories as binary rather than dimensional constructs is limiting Op de Beeck (2025). This might have consequences on our results. For instance the correlation between RT and activation from PPA in response to images of scenes (see Fig. 2B) might be positive in the absence of the lowest activation outlier (see Fig. S8), which contains bodies. Without the forced categorisation of this image as a scene, behaviour and brain might have been positively correlated, as is well captured in the ratings - ROI correlations, which shows that PPA is inversely correlated to how scene-like images are, while this metric is strongly correlated with RTs.

Finally, our experiment was not designed to directly address the nature of the downstream readout that could be operated from OTC activation. As such, we can only make limited guesses on how exactly the activity in our ROIs could be used to guide behaviour. First, we report that it could be linearly readout, but we cannot exclude a non-linear mapping with downstream areas. Additionally, we cannot arbitrate whether a single decoder or several decoders would be best fitted to extract meaningful information from this readout.

## 4.5 Conclusion

Using a DTB approach, we showed that activity in category-selective patches of cortex are relevant to categorisation behaviour, as evidenced by correlations with categorisation performance. While not all ROIs correlate with images from their preferred category, we found a consistent relationship with non-preferred images, across tasks (basic and superordinate categorisations) and across behavioural metrics (RT and motor movements). Taken together, these results highlight the mechanisms through which category-selective activity in visual cortex can guide categorisation behaviour.

## 5 Data availability statement

Data and code to reproduce the analyses presented in this article are available at [this URL](https://github.com/TimManiquet/cortical_selectivity).

## 6 Author contribution statement

Conceptualisation: TM, HOB; data curation: TM, HF, NARM; formal analysis: TM, HF; funding acquisition: TM, HOB; investigation: TM, HF, NARM; methodology: TM, HOB; project administration: HOB; resources: NARM, HOB; software: TM; supervision: HOB; validation: HOB; visualisation: TM; writing - original draft: TM; writing: review & editing: TM, HOB, NARM.

## 8 Diversity in citation practices

The authors of this article report its proportions of citations by gender category to be as follows: M/M = .655; W/M = .207; M/ W = .103; W/ W = .034.

