## Supplementary materials for "Can category-selective cortex predict categorisation behaviour?"

#### 1 Activation-ranked image-wise ROI activation and RT

The figures below (Figs. S1 - S4) reproduce plots from figure 2B with different details. ROI activation is plotted for each image, ranked from the highest to the lowest activation (left y axis, bars). RT for each image is superimposed (right y axis, dots).

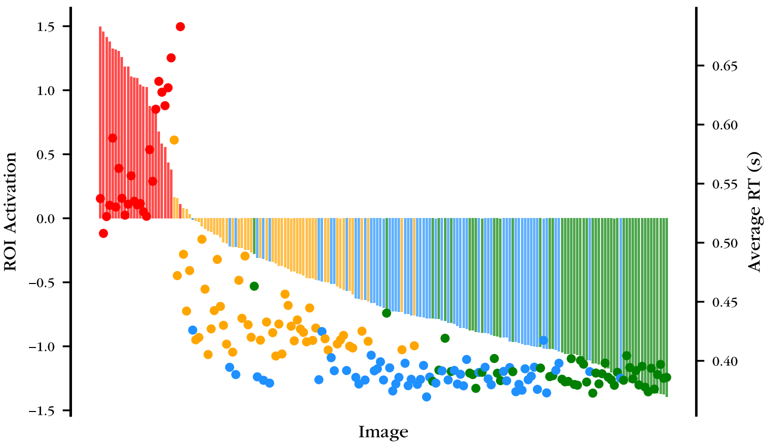


Figure S1: Activation-ranked ROI activation and RT per image for **FFA**. For each stimulus image, a bar represents the mean standardised activity of FFA (left y axis), and a dot represent the mean RT in categorising this image as a face/non face.

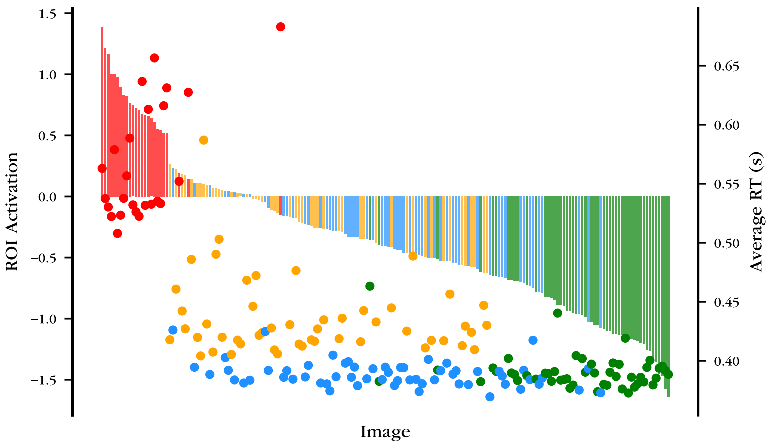


Figure S2: Activation-ranked ROI activation and RT per image for **OFA**. For each stimulus image, a bar represents the mean standardised activity of OFA (left y axis), and a dot represent the mean RT in categorising this image as a face/non face.

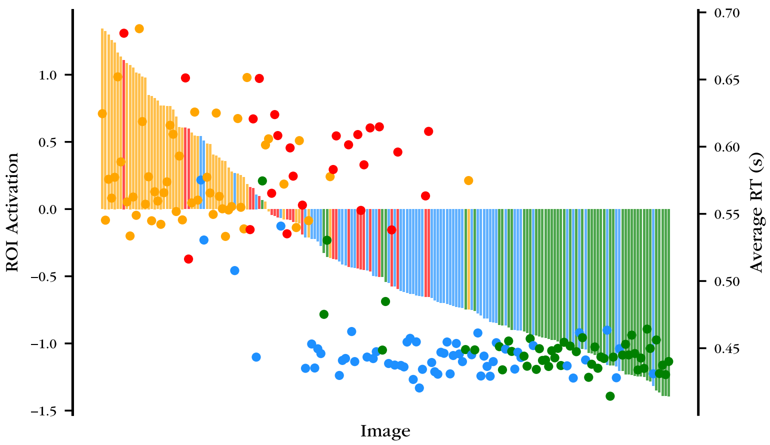


Figure S3: Activation-ranked ROI activation and RT per image for **EBA**. For each stimulus image, a bar represents the mean standardised activity of EBA (left y axis), and a dot represent the mean RT in categorising this image as a body/non body.

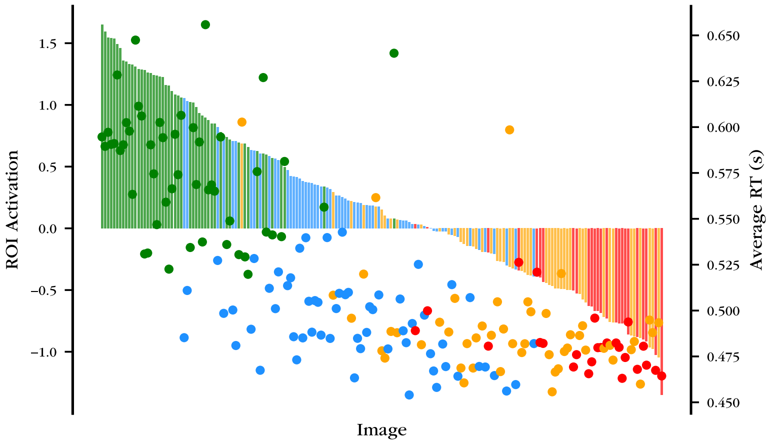


Figure S4: Activation-ranked ROI activation and RT per image for **PPA**. For each stimulus image, a bar represents the mean standardised activity of PPA (left y axis), and a dot represent the mean RT in categorising this image as a scene/non scene.

#### 2 Preferred images per ROI

Figures S5 to S8 display the 5 highest and 5 lowest activation images for each ROI. The selection was made by ranking images per mean ROI activation and picking the top and bottom 5 images, for each category.

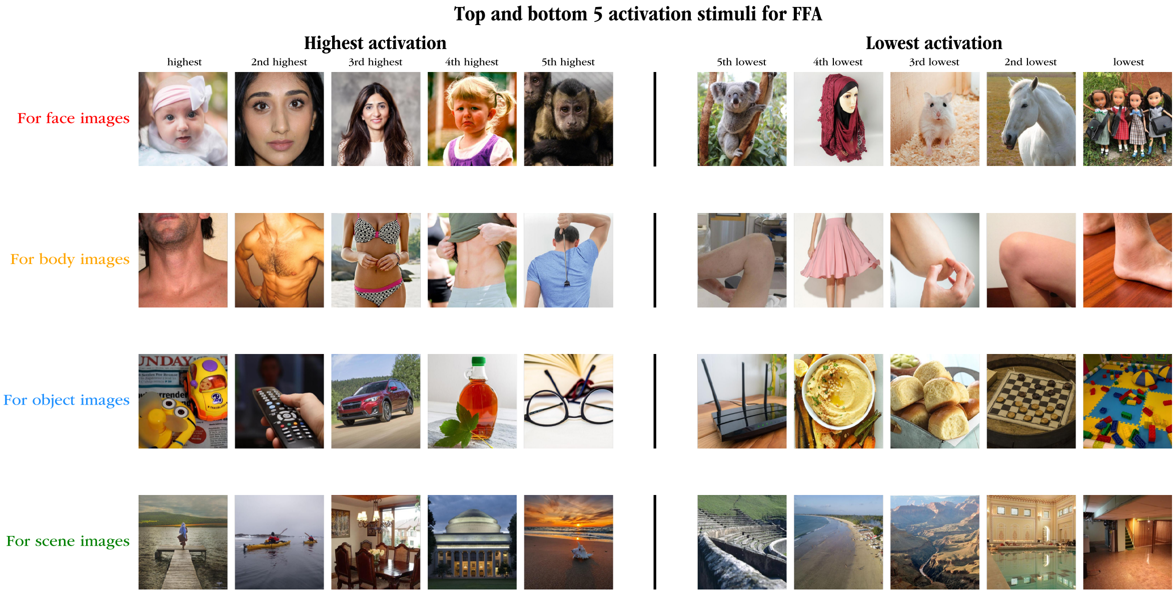


Figure S5: Favourite and least favourite images for **FFA**. Per category of images, the top 5 images with the highest (left) and lowest (right) activation.

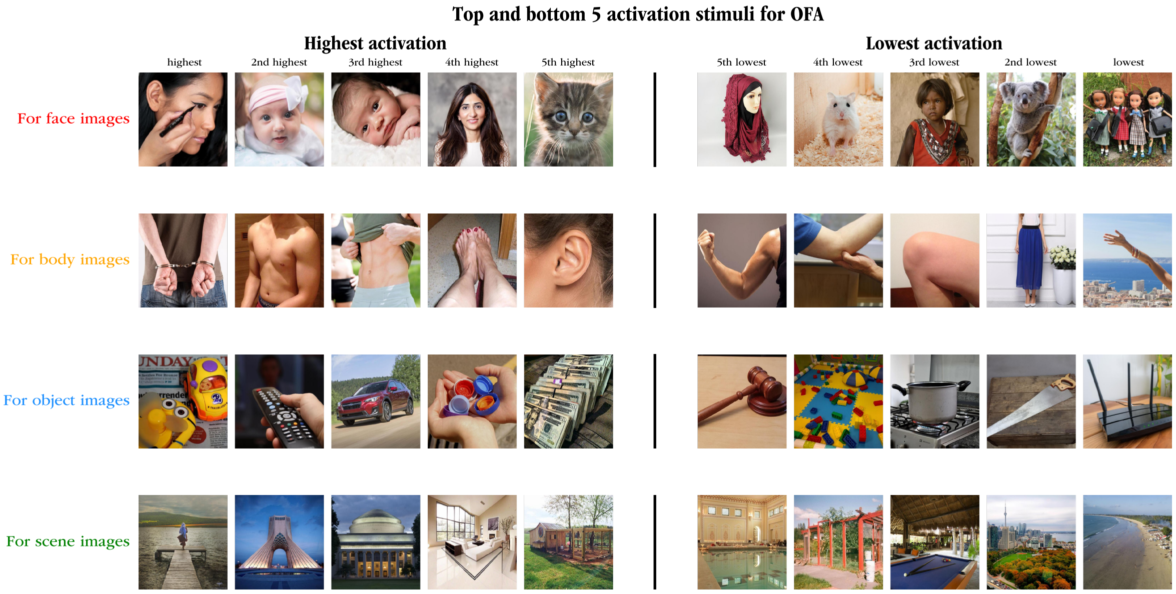


Figure S6: Favourite and least favourite images for **OFA**. Per category of images, the top 5 images with the highest (left) and lowest (right) activation.

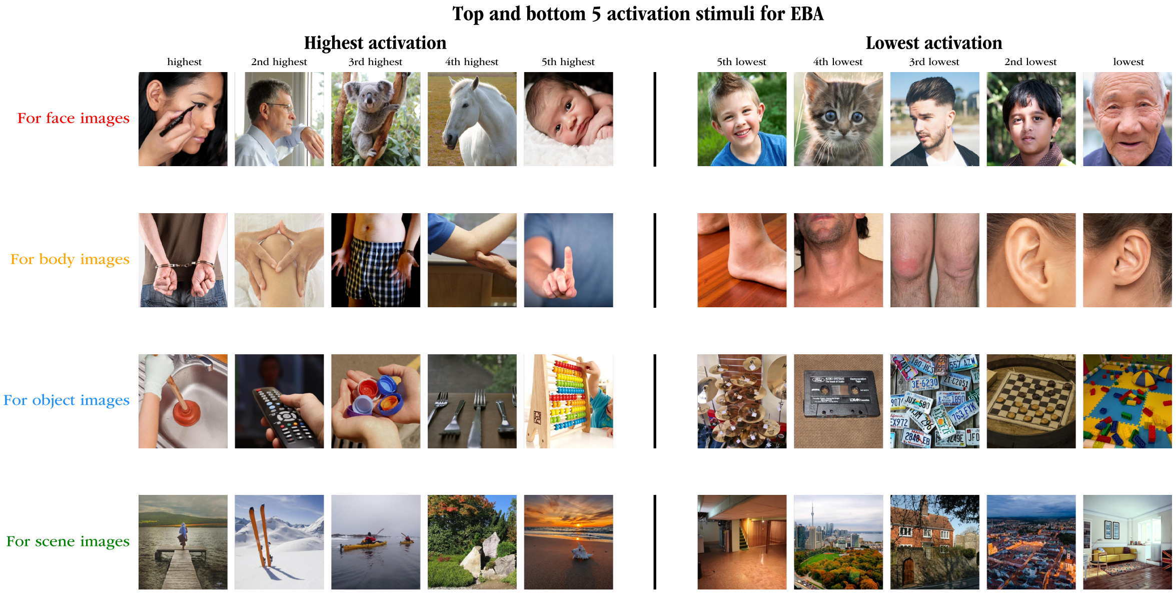


Figure S7: Favourite and least favourite images for **EBA**. Per category of images, the top 5 images with the highest (left) and lowest (right) activation.

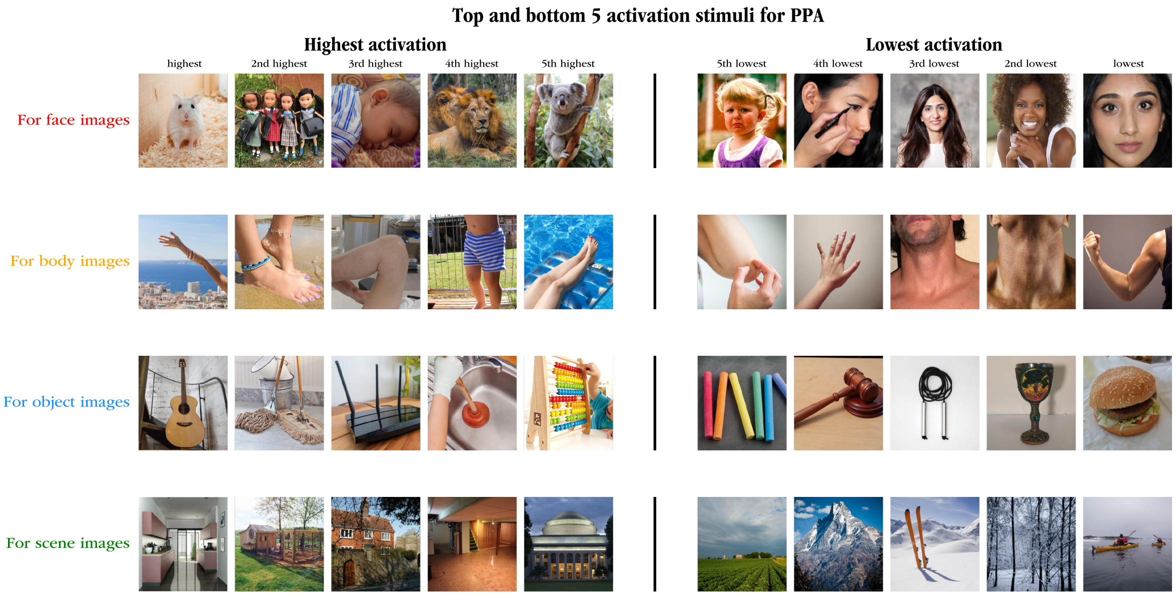


Figure S8: Favourite and least favourite images for **PPA**. Per category of images, the top 5 images with the highest (left) and lowest (right) activation.

### 3 Fastest and slowest images in the categorisation task

Figures S9 to S11 display the top 5 images with the fastest and slowest average RT in the basic categorisation task (see Fig. 2). For each target category (*is this a **face**? yes/no, is this a **body**? yes/no, etc.*), we ranked images from each category by their average RT, and picked the top and bottom 5 images to show.

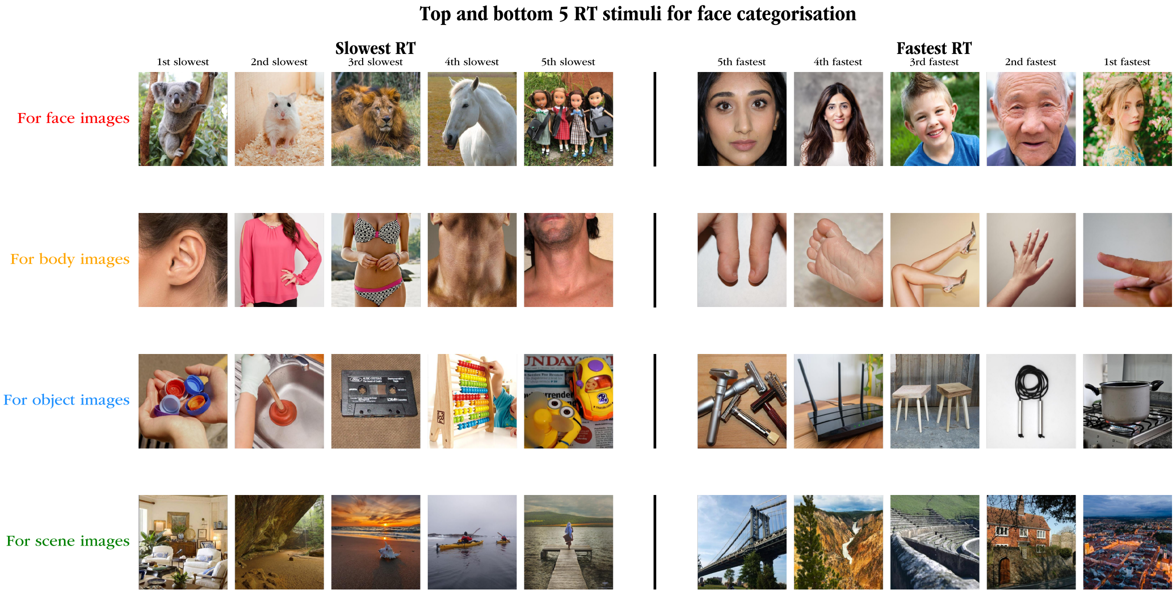


Figure S9: Fastest and slowest images per category for the **face** categorisation task.

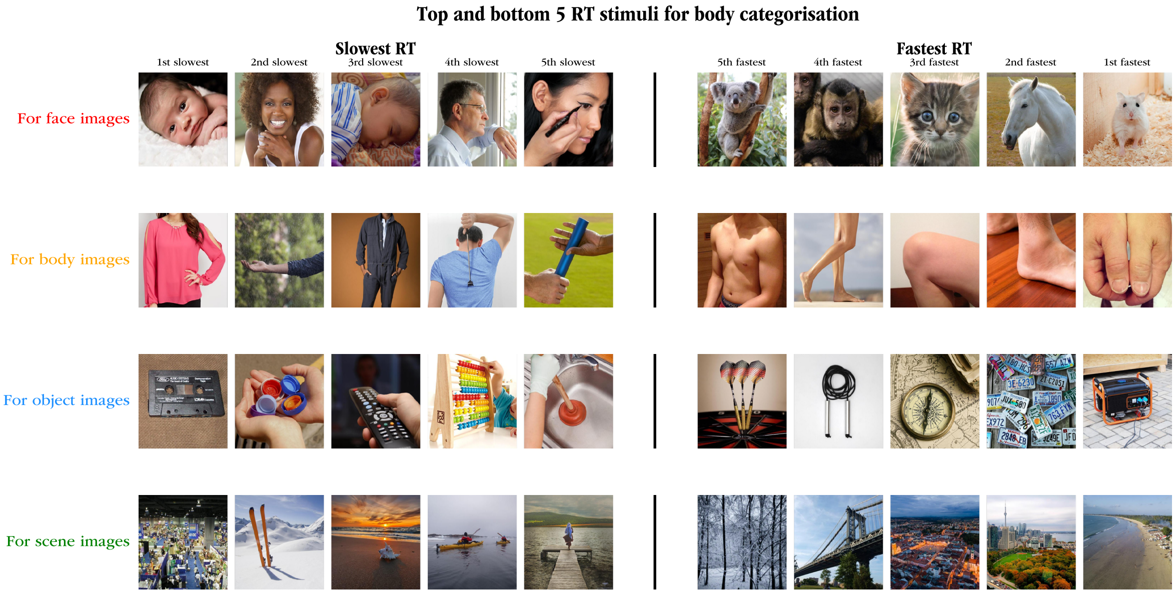


Figure S10: Fastest and slowest images per category for the **body** categorisation task.

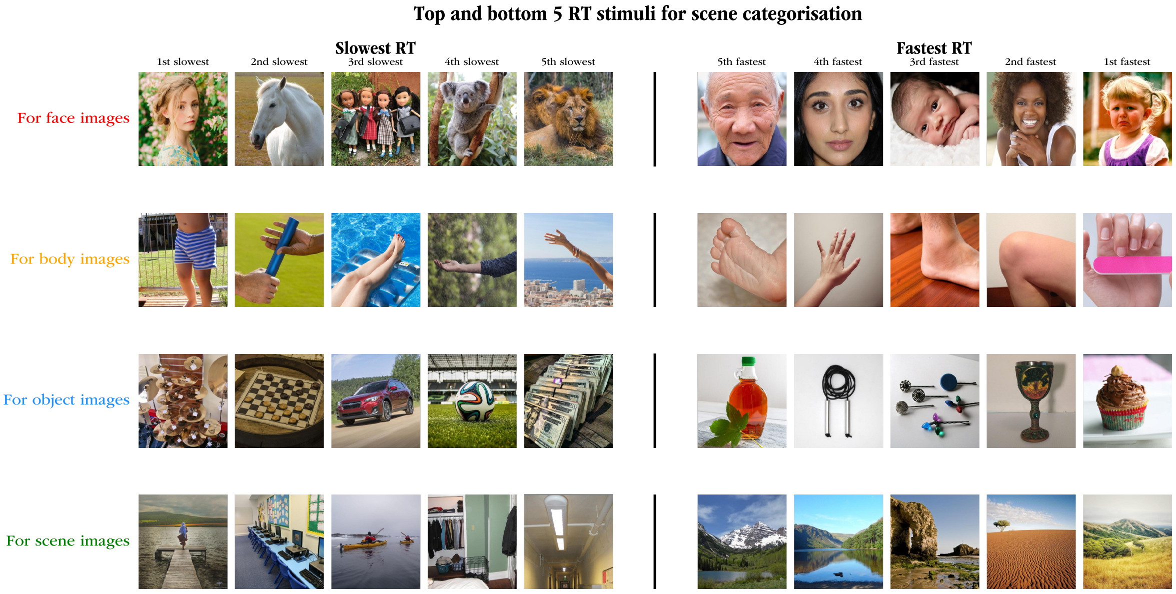


Figure S11: Fastest and slowest images per category for the **scene** categorisation task.

### 4 Largest and smallest area under the curve for motor tasks

In this section, we replicate the top/bottom image analysis from Figures S9 to S11, taking behavioural performance from the motor tasks. Since motor movements extend in time, one could find the top and bottom images based on their horizontal position, creating one series of top and bottom images per time point. However, this plotting obviously be cumbersome and difficult to read. To find the top and bottom images for categorisation tasks using motor movements, we therefore calculate the area under the curve (AUC) per image.

Figures S12 to S15 show the result of this approach, both for basic categorisation and superordinate categorisation. For each categorisation task done with motor movements, we calculated the AUC under the average curve of each image, resulting in a single score that summarises how strongly and quickly each image drifts towards its correct answer button. From this metric, one can expect to find effects similar to those of average RT per image: images that take a long time to be categorised will drift slower, and hence have a smaller AUC. Conversely, images that are quick to recognise will move quickly to the correct answer button, and hence have a larger AUC. As a result, large AUC images in the figures below corresponds to fast RT images in figures S9 to S11.

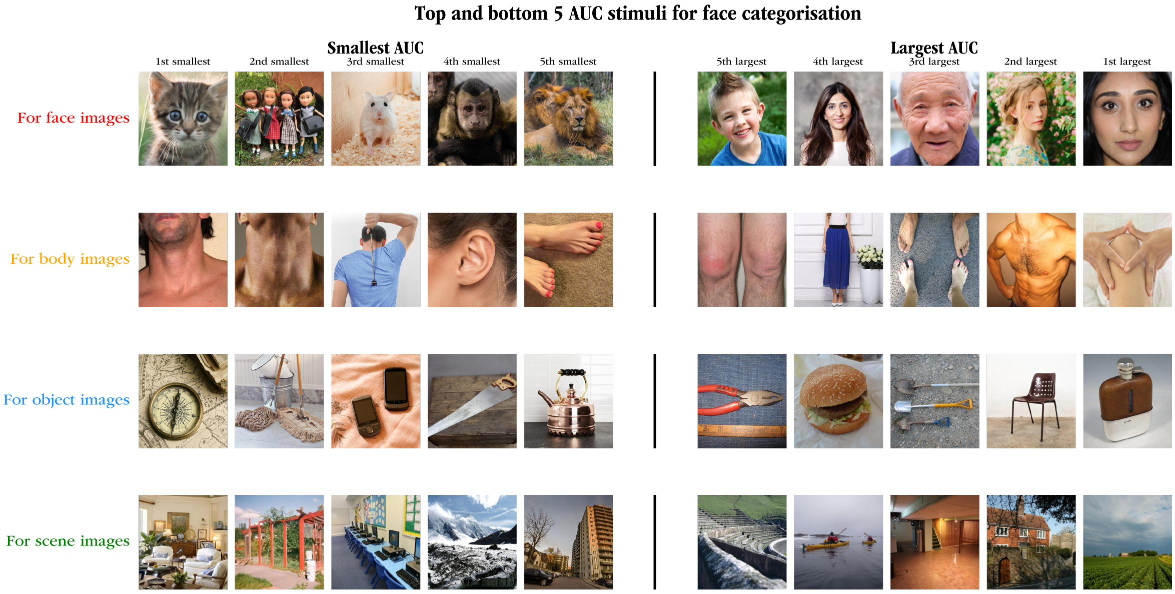


Figure S12: Largest and smallest AUC per image and per category for the **face** basic motor categorisation task.

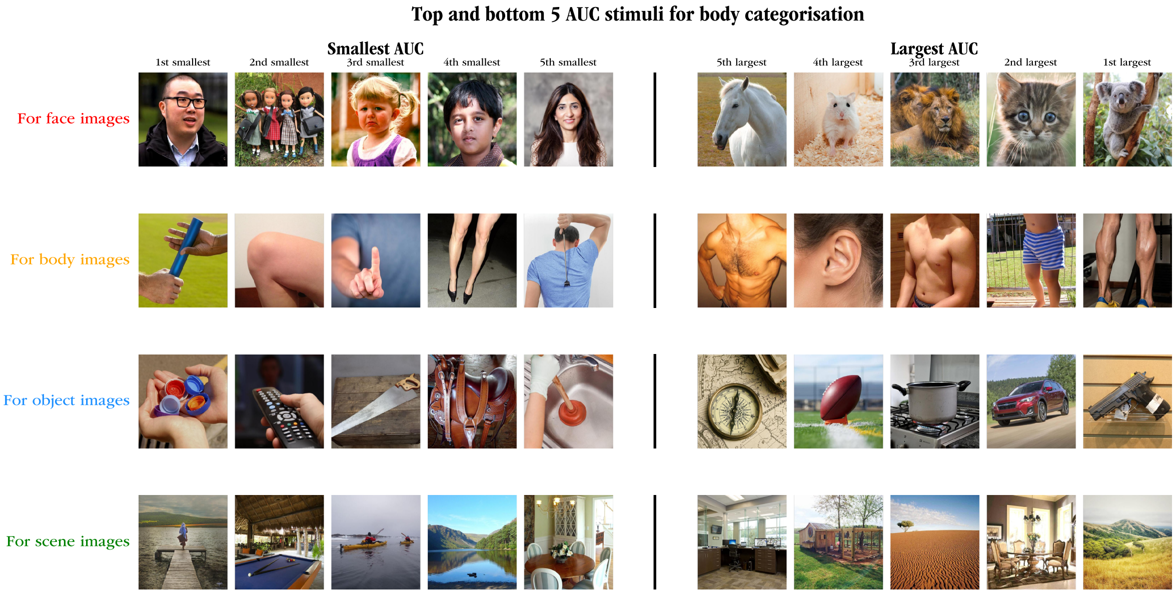


Figure S13: Largest and smallest AUC per image and per category for the **body** basic motor categorisation task.

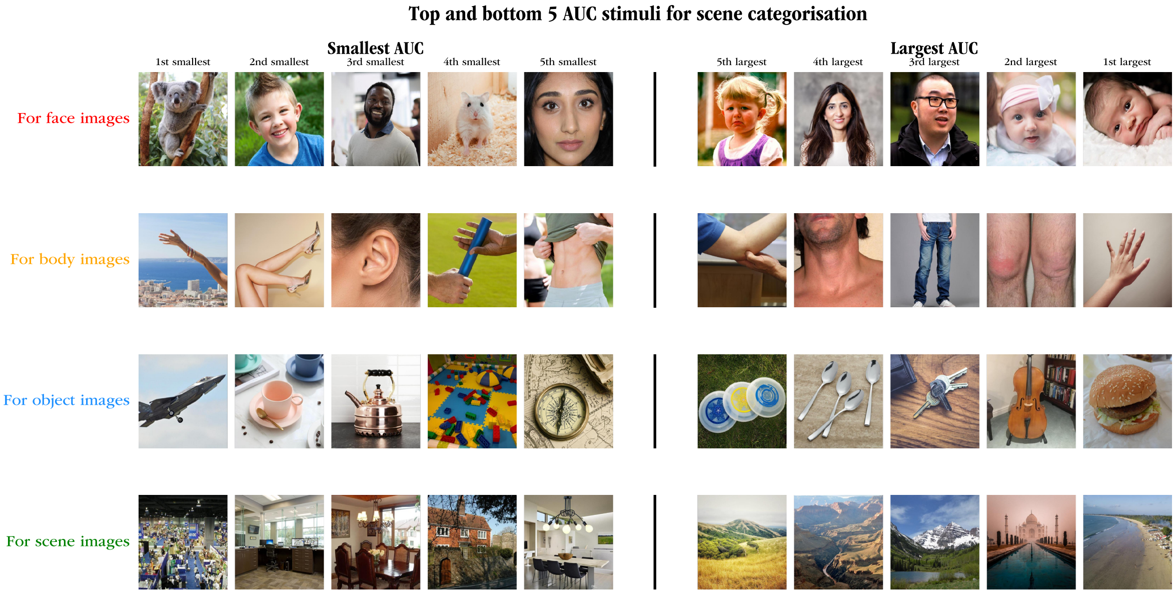


Figure S14: Largest and smallest AUC per image and per category for the **scene** basic motor categorisation task.

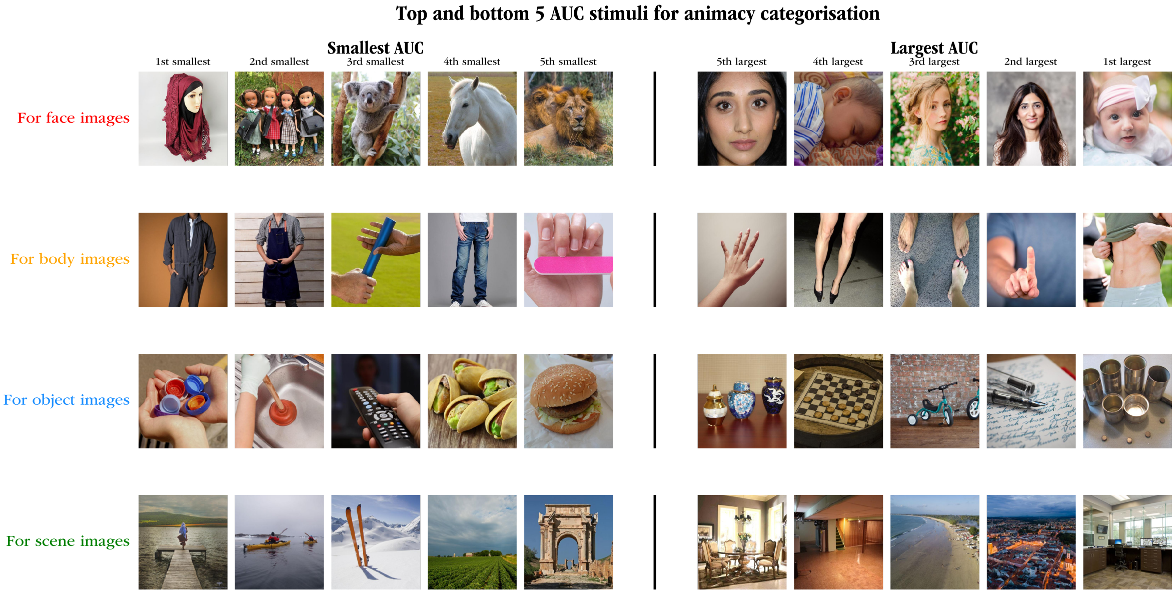


Figure S15: Largest and smallest AUC per image and per category for the super-ordinate **animacy** motor categorisation task.
